# TMELand: An end-to-end pipeline for quantification and visualization of Waddington’s epigenetic landscape based on gene regulatory network

**DOI:** 10.1101/2023.06.07.543805

**Authors:** Lin Zhu, Xin Kang, Chunhe Li, Jie Zheng

**Author notes:** (Corresponding authors: Jie Zheng and Chunhe Li.).,. The first two authors contributed equally to this work.

## Abstract

Waddington’s epigenetic landscape is a framework depicting the processes of cell differentiation and reprogramming under the control of a gene regulatory network (GRN). Traditional model-driven methods for landscape quantification focus on the Boolean network or differential equation-based models of GRN, which need sophisticated prior knowledge and hence hamper their practical applications. To resolve this problem, we combine data-driven methods for inferring GRNs from gene expression data with model-driven approach to the landscape mapping. Specifically, we build an end-to-end pipeline to link data-driven and model-driven methods and develop a software tool named TMELand for GRN inference, visualizing Waddington’s epigenetic landscape, and calculating state transition paths between attractors to uncover the intrinsic mechanism of cellular transition dynamics. By integrating GRN inference from real transcriptomic data with landscape modeling, TMELand can facilitate studies of computational systems biology, such as predicting cellular states and visualizing the dynamical trends of cell fate determination and transition dynamics from single-cell transcriptomic data. The source code of TMELand, a user manual, and model files of case studies can be downloaded freely from https://github.com/JieZheng-ShanghaiTech/TMELand.

## 1 Introduction

WADDINGTON’S epigenetic landscape was proposed by C. H. Waddington in 1957, as a metaphor describing the developmental process of cell differentiation [1]. In his theory, a marble rolling down a hill into a stable locally lowest point (i.e., an attractor) represents a cell differentiating into a specific cell type, and this process is regulated by underlying gene regulatory networks (GRNs). Therefore, different cell types can be identified as different attractors on the landscape. The landscape concept has been extended to studies of reverse differentiation processes such as stem cell reprogramming [2], [3]. The quantification of the epigenetic landscape has become a powerful tool for analyzing cell fate determination [3], [4], [5], [6], [7], [8] or progression of diseases (such as cancer) [9], [10], [11], [12], [13], [14], [15]. There are mainly two types of methods for landscape quantification: data-driven and model-driven. The data-driven methods include those based on Hopfield network [6], [8], [9], [14] and entropy-based methods [16], [17]. The model-driven methods take models of GRN as input, including discrete Boolean network (BN)-based models [5], [18], [19] and continuous differential equation (DE)based models [3], [4], [20], [21].

With the rapid development of single-cell technology, large amounts of scRNA-seq data have been released. To extract system-level information from these data, we aim to use Waddington’s landscape to reveal the cellular dynamics driven by GRNs. However, the data-driven methods for landscape modeling directly infer landscapes from data, without modeling the underlying GRNs explicitly. Thus, compared with the model-driven methods, the data-driven methods lack interpretability to some extent. Here, we propose a pipeline that integrates data-driven algorithms for inferring GRNs from single-cell data and model-driven methods for landscape modeling. Moreover, since most algorithms for constructing landscapes are deployed in the form of source code, users need to adapt the code to run it, which might be a high barrier for researchers with little experience in programming and hamper the popularization of the algorithms. In this case, a user-friendly software tool is desirable. Recently, two software tools with graphical user interfaces (GUIs) have been developed to quantify and visualize Waddington’s landscape, i.e., NetLand [22] and ATLANTIS [23]. NetLand is a Java program for quantifying and visualizing Waddington’s landscape based on DE models of GRN, of which the ideas and code were developed by the corresponding authors of this paper and their colleagues [3], [22]. ATLANTIS is a MATLAB toolbox that visualizes landscapes based on BN models of GRN. In addition to the static landscape, state transition paths can reflect cellular dynamics represented by the temporal evolution of gene expression levels regulated by GRNs. However, NetLand doesn’t offer the functionality of finding the state transition paths. ATLANTIS extracts paths from an initial state to a terminal steady state without mapping the continuous surface of a landscape [23], and thus lacks sufficient details for understanding the dynamical processes along the paths. We develop an open-source, cross-platform Python application named TMELand, based on the truncated moment equations (TME) method for landscape construction and minimizing transition action for calculating state transition paths previously developed by ourselves [20]. Compared with NetLand, TMELand uses an adapted algorithm that can construct a more precise landscape. Meanwhile, compared with ATLANTIS, TMELand can provide more detailed information along the transition paths which can reveal intermediate states, such as partially reprogrammed stem cells. Furthermore, by integrating single-cell data, TMELand builds an end-to-end pipeline, which can make it more widely used compared with tools for either GRN modeling or landscape modeling only. We compare ATLANTIS, NetLand, and TMELand in terms of programming languages, input data, and functionalities, etc. in Table 1.

**TABLE 1.**
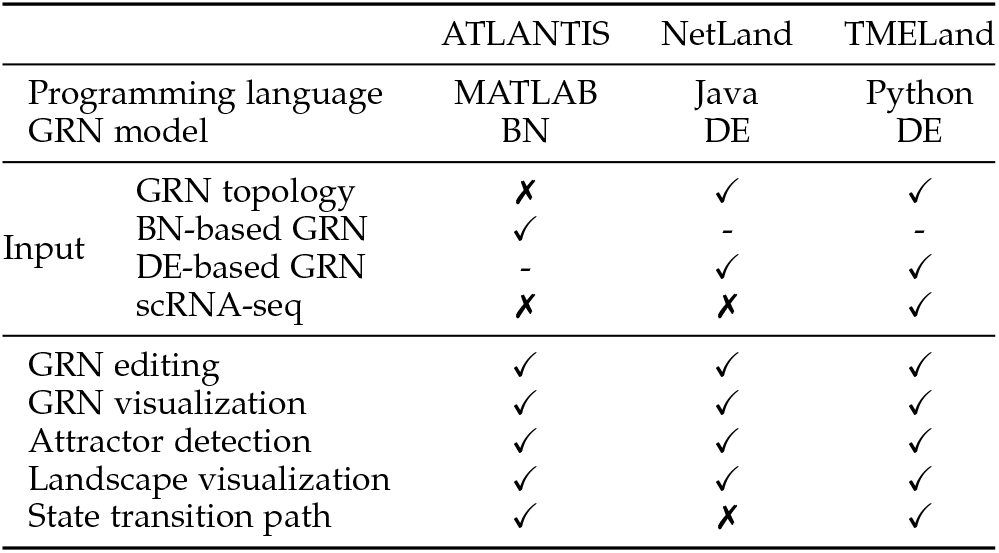
The comparison among ATLANTIS, NetLand, and TMELand

TMELand accepts three types of input data. The first two types are GRN topology in TSV file format and DE-based GRN in the file format of XPPAUT ODE [24] or Systems Biology Markup Language (SBML) [25]. Moreover, TMELand is an end-to-end pipeline which allows single-cell gene expression data (e.g., scRNA-seq) as the third kind of input. As illustrated in Fig. 1, the end-to-end pipeline of TMELand is described as follows. Given the input of a single-cell gene expression matrix and corresponding time series data, we integrate a functionality of visual analytics of the single-cell data and an algorithm for inferring GRN topology from the input data. Users can manually edit the inferred GRN to improve its reliability. With an inferred GRN topology and user-defined parameters, TMELand models the dynamics of the GRN (in non-linear differential equations), simulates trajectories and constructs the corresponding epigenetic landscape, which is visualized in a 3D space. To further study the landscape, users can draw state transition paths by choosing two attractors as the source and target, which helps users map cell types on the landscape and elucidate the order of genes switching on and off in the process of cell fate determination. Note that if the input is a DE-based GRN model, TMELand can parse and extract ODEs (ordinary differential equations) to formulate the cellular dynamics directly.

**Fig. 1.**
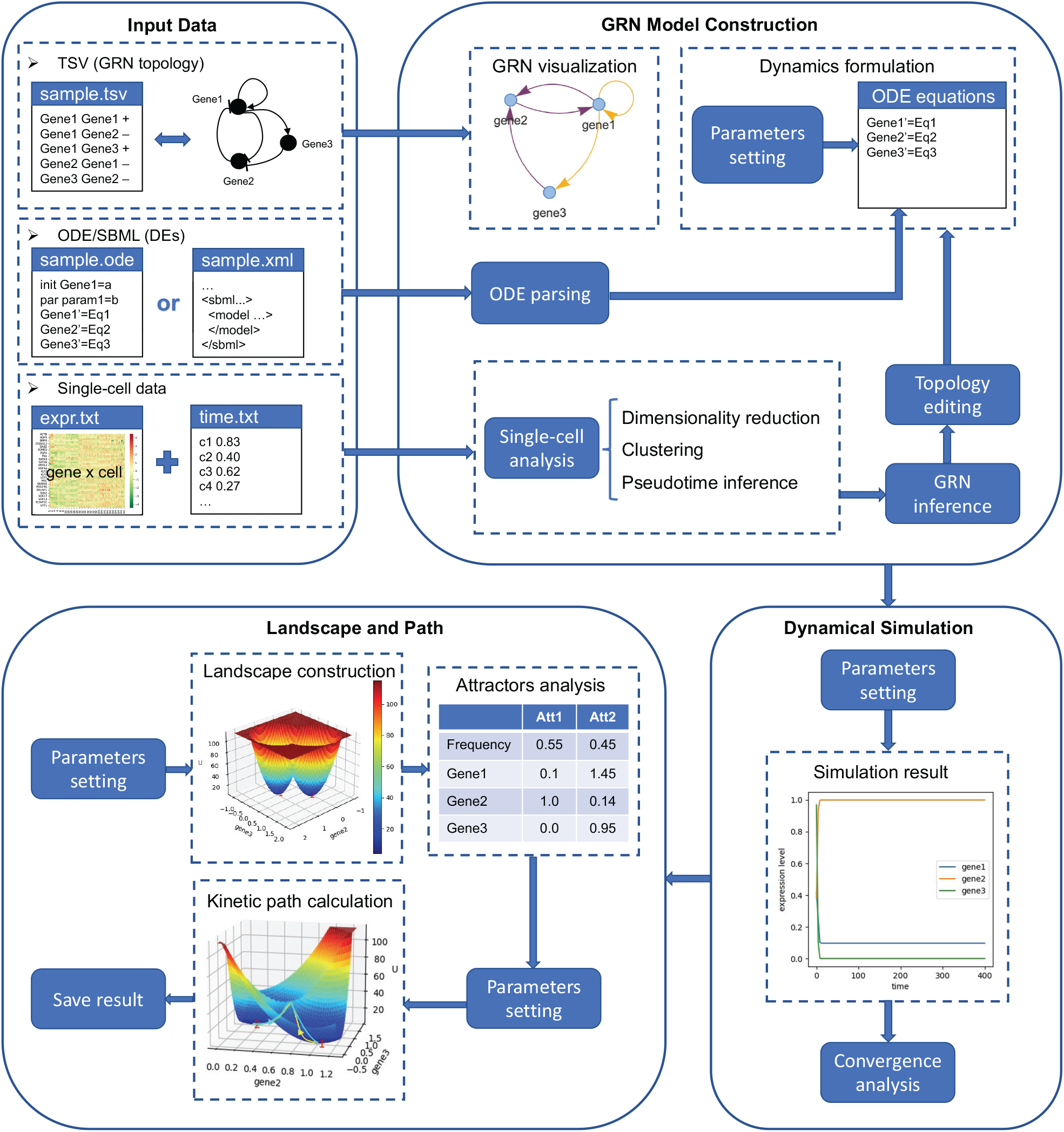
The workflow of TMELand. Input Data: The input data can be one of three types: GRN topology, DE-based model of GRN, and singlecell gene expression data. GRN Model Construction: For the three types of data, there are several modules of data processing in the GRN model construction stage, including GRN visualization, ODE parsing, single-cell data analysis, GRN inference, topology editing, and dynamics formulation. Dynamical Simulation: The dynamical simulation is used to find appropriate parameters to find stable states. Landscape and Path: The final stage consists of the calculation and visualization of the epigenetic landscape and kinetic paths.

In this paper, our contributions are summarized as follows:

- We propose an end-to-end pipeline integrating datadriven GRN inference and model-driven landscape mapping, including single-cell data analysis, GRN topology inference, and landscape quantification and visualization.
- We implement a state-of-the-art algorithm [20] modeling Waddington’s epigenetic landscape and calculating state transition paths, and encapsulate it as the core part of a user-friendly Python software named TMELand.

## 2 Related Work

This section will give a brief introduction to epigenetic landscape modeling and state transition path calculation. As mentioned above, existing methods for modeling Waddington’s epigenetic landscape can be divided into two categories: data-driven and model-driven methods.

### 2.1 Data-driven Methods

Data-driven methods typically reconstruct landscapes from gene expression data, including Hopfield network-based methods and entropy-based methods. Hopfield network, a classic type of artificial neural network, is an associative memory model, which can store input patterns as attractors in the network and the patterns can be recalled from partial input [26]. So far, most of the data-driven methods use the Hopfield network to model the epigenetic landscape. Lang et al. used a Hopfield network to analyze stem cell reprogramming experiments, and their model was able to predict reprogramming protocols and key reprogramming factors consistent with known experimental results [6]. Ragan et al. modeled landscapes based on a discrete Hopfield network and applied it to studies of the cellular development and disease progression [7], [9], [12]. Guo et al. used a continuous Hopfield network to map landscapes from singlecell gene expression data and infer pseudotimes of cells on the landscape [8]. Conforte et al. constructed an attractor landscape based on Hopfield networks to analyze breast cancer data [14].

Another type of methods for quantifying landscapes is based on entropy, which quantifies the promiscuity of a cell state to the elevation of each point in a landscape. With the decrease of elevation (i.e., entropy reduction), cells tend to differentiate into stable cell types. Banerji et al. used the concept of network entropy to estimate signaling pathway promiscuity and they demonstrated that network entropy can quantitatively measure the cell state of differentiation thereby defining the elevation of the Waddington’s landscape [16]. Shi et al. assessed four methods (including a network entropy-based method) for measuring singlecell potency using data from nine independent scRNA-seq experiments, which revealed that the robustness of singlecell potency measurement is driven by correlations between mRNA expression and network connectivity [17].

### 2.2 Model-driven Methods

Model-driven methods take GRN models as input, such as discrete BN models and continuous DE models. A BN model of GRN is encoded on a directed graph, where each node corresponds to a gene and contains a binary variable representing the gene’s expression level, and each directed edge represents the regulatory interaction between two genes. The gene expression of a node is updated by a Boolean function consisting of logical operators. AlvarezBuylla and Davila-Velderrain used a BN model of GRN to study flower development, and projected cells of flower tissue to attractors [18], [19]. Flöttmann et al. constructed a probabilistic BN and visualized the probability distribution of states at time scale as an epigenetic landscape [5]. Cho et al. modeled an attractor landscape based on a BN to analyze colorectal cancer progression and therapies [10], [11], [13].

However, as a discrete model, BN tends to oversimplify different gene expression levels and time scales, which could lead to loss of crucial information. By contrast, continuous DE models can provide more refined dynamical information. A DE-based GRN model comprises a set of differential equations that describe the dynamics of transcriptional regulation, e.g., the temporal evolution of gene product concentrations driven by the molecular interactions in the network. Wang et al. modeled a GRN as Hill function-based nonlinear differential equations and proposed the concept of potential landscape [2], [3], [20]. Bhattacharya et al. proposed a method to quantify epigenetic landscapes using deterministic quasi-potential calculated using ODE models of 2-gene GRNs [4]. Recently, Zhang et al. proposed a Monte Carlo-based method to generate landscape by using simulations of DEs in an efficient way [21].

### 2.3 State Transition Path

Based on the attractor landscape, finding the state transition paths between attractors can reveal details of cellular dynamics, such as changes in gene expression and key states in transition. ATLANTIS generates trajectories after constructing landscapes, but it only displays the order of states lacking detailed information such as positions of states and their distances on the landscape. In DE-based model-driven methods, the state transition paths can be found by path integral [27] or minimum action approach [28]. The path integral approach formalizes the probability of the network dynamics from an initial state to end state as an integral, by representing the sum of all possible paths with different weights, and selecting the path with the largest weight as the optimal transition path [27]. The minimum action approach, another method to calculate the most probable transition path between steady states, minimizes the transition action functional over all possible paths [28]. We adopt the minimum action approach as the algorithm for state transition path calculation in TMELand.

## 3 Methods

### 3.1 DE-based GRN

In TMELand, to model DE-based GRNs, we assigned stochastic dynamics to a static GRN with Langevin equations [20]. Given the topological structure of a GRN, we used Equation (1) to describe the temporal evolution of the expression level of each gene. In Equation (1), *x*_*i*_ is the expression level of the *i*th gene, *F* (*x*_*i*_) is the driving force and takes a nonlinear ODE form as shown in Equation (2), and *ζ* is the Gaussian white noise, which is related to the user-defined diffusion coefficient. In Equation (2), *m*_*i*1_ is the number of activators of gene *i, m*_*i*2_ is the number of inhibitors of gene *i*, 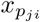 is an activation gene of gene *i*, 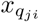 is an inhibition gene of gene *i, A* is the activation constant, *B* is the inhibition constant, *n* is the Hill coefficient, *s* is the threshold, and *k* is the degradation rate. All the aforementioned parameters will be decided by GRN topology and user input.

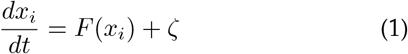

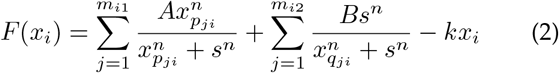

### 3.2 Quantification of Potential Landscape by TME

After constructing a DE-based model of GRN, we used the corresponding Fokker-Planck equation (FPE) to describe the density function. When the diffusion coefficient is small (corresponding to weak noise), the solution of FPE can be approximated by a Gaussian distribution based on the first two moments, i.e., the mean and covariance matrix [20], [29]. By solving the force system (i.e., Equations (2)) numerically, we first acquired the steady-state points 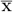 as the mean matrix, also known as attractors. Then, from the calculated steady-state points 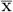 and the diffusion coefficient characterizing the magnitude of noise level, we calculated the covariance matrix *σ*. We quantified the landscape using the weighted sum of multiple normal distributions *P*_*ss*_ with mean 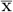, covariance matrix *σ*, and weight vector **w**, where **w** contains the frequency of each stable state. With the steady-state probability distribution *P*_*ss*_, we can calculate the potential landscape as *U* = *−ln*(*P*_*ss*_) [3], [20].

### 3.3 Calculation of Kinetic Transition Paths

After quantifying the potential landscape, TMELand calculates kinetic transition paths among different attractors to depict the dynamic processes of cell state transitions. TMELand adopts the strategy of minimizing transition actions to obtain the minimum action paths (MAPs) as kinetic transition paths [28] using the trust-region constrained optimization algorithm [30].

## 4 Functionalities and Implementation

TMELand was developed using the Python language and its GUI framework, Tkinter, which takes advantage of the open ecology of Python, such as abundant scientific computation and visualization libraries (e.g., NumPy [31], SciPy [32], pandas [33], Matplotlib [34]). Additionally, since many software tools are developed using R in Bioinformatics, we used conda to manage the environment to facilitate the integration of Python and R code. We provided a package list to allow for one-click installation. This section will give a brief description of the functionalities of TMELand, and more details can be found in the User Manual.

### 4.1 Construction of GRN Models

TMELand accepts input in three formats. The first is the GRN topological structure, which reflects the regulatory relationships among genes. The second is the DE-based dynamical model of GRN. The third is single-cell gene expression data, such as scRNA-seq data, from which TMELand uses a GRN inference algorithm to infer network topology.

#### 4.1.1 Network Structure

GRN structure is represented in TSV format which is typically used to represent a directed graph, i.e., node 1 and node 2 followed by a gene regulatory relationship, with ‘+’ denoting activation and ‘-’ denoting inhibition. After the topological structure of a GRN is loaded, it would be visualized on a panel by a click on the ‘GRN topology’ button. The graph visualization tool is Jaal (https://github.com/imohitmayank/jaal), which is launched by a child process, and then it can be accessed from a specified port of the localhost. On the panel, users can search for a node, move the graph, zoom in the display, drag a node, and highlight an edge when the cursor focuses on it, which is especially useful when loading a dense GRN structure (Fig. 2A). Next, to assign dynamical features for the loaded GRN, users can open the ‘DE-based GRN’ window and set values of parameters mentioned in Section 3.1. Then, TMELand generates nonlinear differential equations using user-specified parameters and shows them in the lower part of the window (Fig. 2B).

**Fig. 2.**
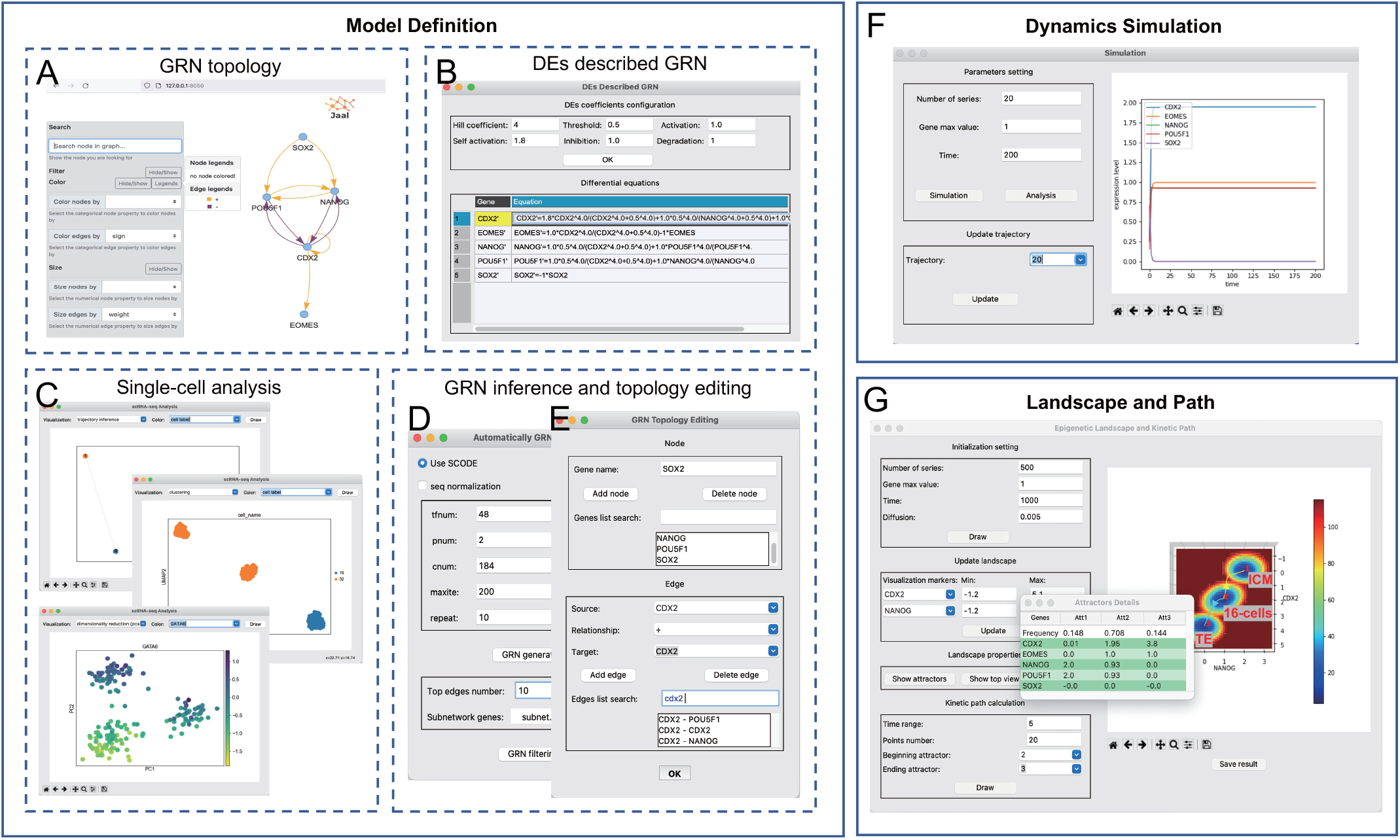
The graphical user interfaces of TMELand using Case Study 1 as an example. A) The editable inferred GRN topology visualization in the browser. B) The dynamic DE modeling for GRN structure. C) The single-cell data visualization and analysis in Case Study 1. D) The user interface of the GRN inference module. E) The GRN topology editing module for the inferred GRN structure. F) Dynamical simulation based on DE models of GRN. G) The visualized landscape, attractors information, and state transition paths for input data.

#### 4.1.2 Differential Equations

DE-based GRN models are described in the XPPAUT ODE file format or SBML format. The imported models will be parsed as a set of differential equations to reflect gene expression regulation dynamics and displayed in the ‘DEbased GRN’ window. TMELand formulates ODEs from an XPPAUT ODE file by its rules and extracts ODEs from an SBML file using libSBML (https://github.com/sbmlteam/libsbml) Python library.

#### 4.1.3 scRNA-seq Data

With the rapid advance of single-cell DNA and RNA sequencing technologies, a large number of single-cell RNA-seq datasets have been published. In order to use these data and further expand TMELand functionalities, we accept scRNA-seq data as an input. TMELand integrates an efficient GRN inference algorithm, SCODE [35], to infer directed and signed GRN topology from scRNA-seq data. When inferring the GRN topology, TMELand sorts the edges by scores and selects top k edges, where k is a user-specified number. Users can also extract a subnetwork by uploading a gene list (Fig. 2D). However, the GRN inference algorithm can’t guarantee that the inferred GRN structure is completely correct. Thus, TMELand supports GRN topology editing. Users can add a node, or delete a node if it is believed not involved in any reaction in the context under study. Users can also add or delete an edge by specifying the source node, regulatory relationship, and the target node. To speed up the node or edge finding process, users can search from their collections (Fig. 2E). Once we obtain the final GRN structure, the rest of the workflow is the same as described in Section 4.1.1. Furthermore, to explore singlecell data, TMELand uses Scanpy [36] to support visual analysis. Specifically, TMELand provides PCA-based linear dimensionality reduction [37], Leiden graph clustering [38], and PAGA-based trajectory inference [39]. Users can highlight a part of the plot by a specific gene, cell names, or clustering results (Fig. 2C).

### 4.2 Dynamical Simulation

In landscape quantification, a crucial step is simulating many trajectories to find steady states. As the convergence of the trajectories directly determines the locations of steady states, users can execute the dynamical simulation of a GRN by solving the associated DEs before mapping the corresponding landscape. With the user-specified number of trajectories, random initial values of gene expression levels within a range, and a time interval, the solutions of the DEs will be displayed as time series on the right panel of the ‘Simulation’ window (Fig. 2F). The ODE solver used in TMELand is the odeint function in the SciPy library of Python, which was developed based on the lsoda function from the FORTRAN library ODEPACK [40]. Users can watch the animated process of convergence of the trajectories, and can additionally use the ‘Analysis’ functionality to analyze the convergence. Also, users can switch trajectories to update the plot.

### 4.3 Landscape Quantification and Analysis

From the simulated trajectories and a user-specified diffusion coefficient, TMELand quantifies and visualizes the corresponding landscape using the aforementioned TME algorithm. The top box in the left frame (shown in Fig. 2G) allows users to set initial parameters of the landscape (e.g. the number of trajectories and the maximum value of gene expression level). The second box allows users to select two marker genes for the x-axis and y-axis respectively and set the value ranges of the expression levels of the two marker genes. The third box shows more properties of the landscape, e.g. the ‘Show attractors’ functionality shows the positions and widths of attractors. To save the information of the constructed landscape, users can click the ‘Save result’ button to save it as a JSON file, which can be reloaded and restored later on for further analysis.

### 4.4 State Transition Paths

Similar to the landscape, to draw the state transition paths, we first reduce dimensionality from multiple genes to the same two marker genes as used for plotting the landscape. In this way, we can obtain the 2D coordinates of the paths found by using the aforementioned algorithm for finding MAPs and project them onto the surface of the landscape. Users can specify path-related parameters (e.g. time range and granularity) and choose the beginning attractor and ending attractor to draw the transition paths. Fig. 2G contains an illustration of the transition paths drawn on the landscape, which exhibits the temporal evolution of the marker genes’ expression values during different dynamical processes. The visualization of state transition paths can yield deeper insights into the dynamics of gene regulatory networks underlying cell fate decisions, such as revealing the driving force behind the induced pluripotent stem cell reprogramming.

## 5 Results

## 5.1 SCA and TME

The self-consistent mean field approximation (SCA) method for landscape quantification [3] used in NetLand is based on an assumption of weak correlations among gene variables (i.e. gene expression levels), which may not be realistic for some situations where the gene variables have strong correlations with each other. By contrast, the TME method [20] explicitly considers the strong correlations among different genes when calculating the steady states *P*_*ss*_. Compared to NetLand, TMELand constructs landscapes that are closer to results from benchmark numerical simulations, meaning that TMELand can construct a more precise landscape than NetLand [20]. Fig. 3 is an example that shows the negative correlation between *GATA6* and *NANOG* in the result from TMELand which is not shown in the result from NetLand for a 52-gene stem cell regulatory network with the same parameters as in the original papers [3], [41].

**Fig. 3.**
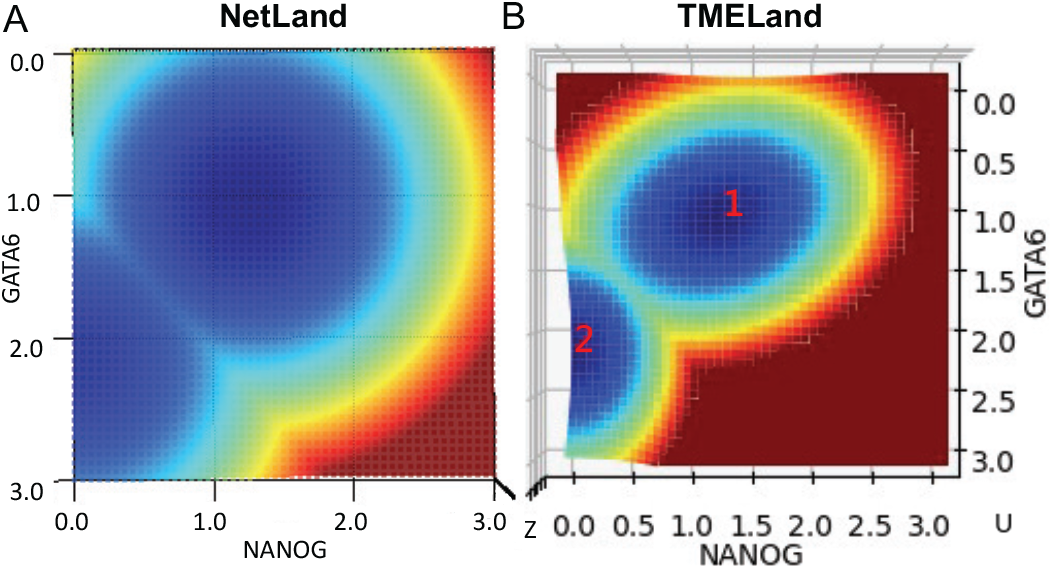
The comparison of landscapes plotted by NetLand and TMELand on the 52-gene stem cell regulatory network [3], [41].

### 5.2 Case Study 1: Early mouse embryonic development

In this case study, we used the single-cell gene expression data from [42] to study the early mouse embryonic development. During the early development from the 1-cell zygote to the blastocyst, three distinct cell lineages are formed, namely the trophectoderm (TE), the primitive endoderm (PE), and the epiblast (EPI). The EPI and PE cells are differentiated from the inner cell mass (ICM) [42]. To focus on the lineage formation of ICM and TE, we extracted the data of cell samples at the 16-cell and 32-cell stages from the original 437 cells. The processed data consist of 48 genes and 184 cells. We used the single-cell data analysis module of TMELand to analyze the data. The result of clustering highlighted by cell labels is shown in Fig. 2C, from which we can see three distinct clusters, corresponding to the three cell types in the dataset (i.e. 16-cell, ICM and TE). We can also use marker genes to identify specific cell types from the clustering results. Then, we used the aforementioned SCODE algorithm to do GRN inference, for which the parameter setting is shown in Fig. 2D. The inferred relationships involve all input genes. Here, we aim to focus on the genes related with the formation of ICM and TE, i.e., *EOMES, CDX2, POU5F1, NANOG* and *SOX2* [43]. For that, we can upload a list of genes to extract a subnetwork induced by the uploaded genes as the output GRN. We can search the literature to assess the accuracy of the inferred network structure and then edit the GRN using the ‘Topology editing’ module (Fig. 2E). Based on the GRN topology (Fig. 2A), we can construct DEs (Fig. 2B), map a landscape and find the state transition paths (Fig. 2G). It is known that *NANOG* is the marker gene of ICM, and *CDX2* is the marker gene of TE, so we set *NANOG* and *CDX2* as the x and y axes respectively to visualize the landscape. *NANOG* has high expression in 16-cell and ICM, and *CDX2* has high expression in 16-cell and TE [42]. Based on such prior knowledge, we can tell that, in Fig. 2G, attractor 1 represents ICM, attractor 2 represents 16-cell state, and attractor 3 represents TE. We can also plot the transition paths from 16-cell to ICM and TE respectively, corresponding to the two cell developmental processes.

### 5.3 Case Study 2: Cancer attractors

In this case study, we used a 6-gene GRN model proposed in [15] for studying the cancer stem cells (CSC). The structure of GRN is shown in Fig. 4A, with black lines representing activation and red lines representing inhibition. In the original work, the authors identified 4 attractors, i.e., CSC, cancer cells, normal cells, and stem cells. We used TMELand to load the ODE model and draw the landscape and transition paths with marker genes (*P53* and *ZEB*) as shown in Fig. 4B. From the results, we can find 8 attractors, among which attractors 1 and 2 can be treated as one merged attractor representing cancer cells (low *P53* and low *ZEB*), and likewise for attractor pairs (3, 4) as CSC attractor (low *P53* and high *ZEB*), (5, 6) as normal cell attractor (high *P53* and low *ZEB*), and (7, 8) as stem cell attractor (high *P53* and high *ZEB*). The expression level of *P53* represents the degree of cancerization, and that of *ZEB* represents the degree of stemness [15]. With the activation of *P53*, the cell state gradually moves from a CSC attractor to a normal cell attractor. We drew paths among attractors 1, 3, 5, and 7 with the yellow lines representing the transition from the CSC state to the normal cell state and the blue lines representing the transition from the normal cell state to the CSC state, and found that the paths are consistent with the original work [15]. Furthermore, we tested the model using both NetLand [22] and the Monte Carlo method for landscape modeling in [21], and both approaches gave 8 attractors as well, which indicates that these methods (including TMELand) can reveal more details than the original work [15].

**Fig. 4.**
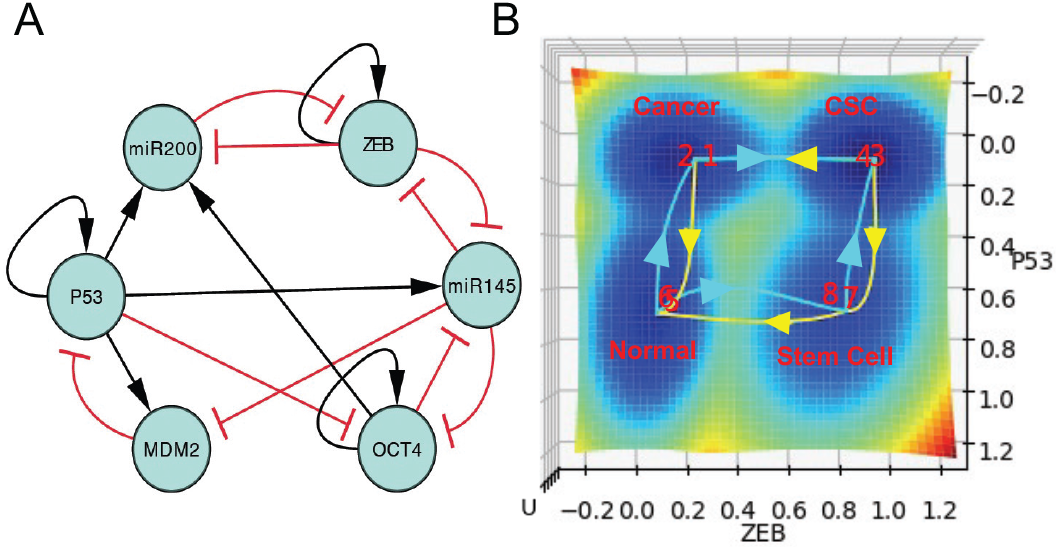
A) The network structure of the 6-gene GRN model [15]. B) The result of Case Study 2 using TMELand.

### 5.4 Comparison of Computational Efficiency

To assess the computational efficiency of TMELand, we compared it with NetLand with the same parameters on three GRN models in SBML format, which describe a 2-gene network [44], a 6-gene network [15] and a 52-gene network [3], [41]. We compared the running time and running peak memory usage between the two software tools, and the average results over 5 runs are shown in Fig. 5. The test was carried out on a laptop computer, with Intel Core i7 and a memory size of 12 GB, under Windows 10.

**Fig. 5.**
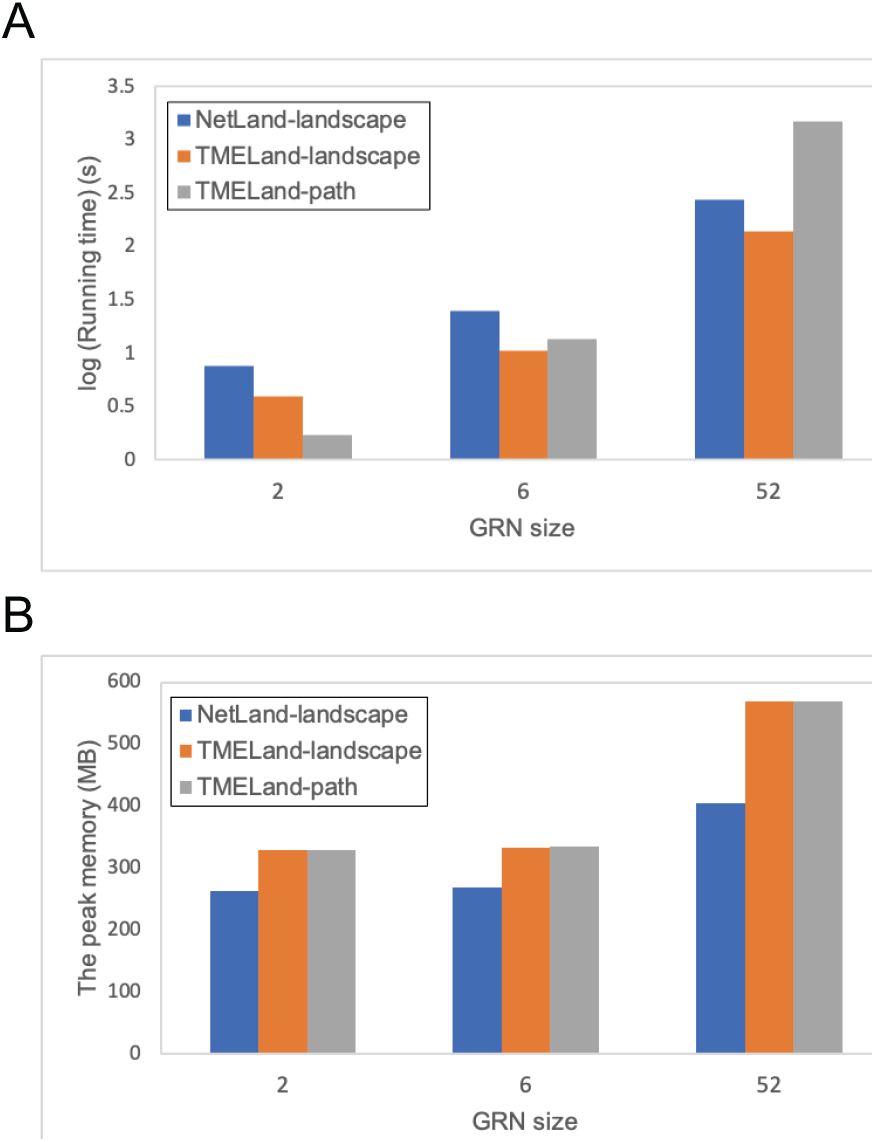
A) and B) are running time and peak memory usage for NetLand and TMELand on three GRN models with different sizes.

Fig. 5A and Fig. 5B show the running time and the memory usage respectively. The blue bars represent the performance of NetLand to calculate the landscape, and the orange and gray bars represent the performance of TMELand to calculate the landscape and paths respectively. Since NetLand cannot calculate the paths, we only recorded its performance on mapping the landscape. From Fig. 5A, we can see that TMELand is faster than NetLand in calculating the landscapes on all the three models, and both of the blue and orange bars grow slowly with network size. However, the running time for calculating the paths grows fast with the network size. From Fig. 5B, we notice that the peak memory usage of TMELand is larger than NetLand when calculating the landscape, but TMELand does not need more memory for finding the paths than quantifying the landscapes in contrast to the running time.

## 6 Conclusion

In this work, we developed an end-to-end pipeline integrating a data-driven GRN inference algorithm with a model-driven method for landscape modeling. It has been implemented into a user-friendly Python software tool named TMELand. Compared with existing data-driven methods for landscape modeling, TMELand uncovers gene regulatory networks underlying the landscapes explicitly, which has better interpretability. Compared with existing model-driven methods for landscape modeling, we automate the process of model construction by integrating a GRN inference algorithm, which lowers the requirements for prior knowledge about GRN models as input. TMELand accepts three types of data as input, i.e., GRN topology, DE-based GRN, and single-cell gene expression data. It then formulates a dynamical model of the GRN, quantifies and visualizes the corresponding Waddington’s epigenetic landscape, and draws state transition paths on the landscape. Compared to our previous software of NetLand, TMELand adopts a more precise algorithm to build landscapes and provides the additional functionality of calculating state transition paths between attractors, which helps reveal cellular dynamics on the landscape more clearly than NetLand.

## 7 Discussion

In the area of modeling the Waddington’s epigenetic landscape, model-driven methods have been widely used thanks to their interpretability for the underlying dynamics of gene regulation. However, it raises a barrier for beginners to build GRN models as input, and it is also limited to smallscale GRN models. In this work, we incorporate a datadriven method to help users build GRN models thereby implementing an end-to-end pipeline from single-cell gene expression data to the visualization of landscape. To the best of our knowledge, GRN inference from gene expression data and modeling of Waddington’s epigenetic landscape, two important problems in Computational Systems Biology, have mostly been studied separately. Here we propose to assemble them into one pipeline, and we argue that this move would benefit not only users of the software tools but also researchers in both areas. However, there are still limitations in TMELand and some extensions can be considered in future work. One is to incorporate more data-driven GRN inference algorithms for users to choose from. Another strategy is to add links from public databases of gene regulatory network models to help users get reliable GRNs. For landscape modeling, a difficulty is that the landscape is hard to evaluate quantitatively, and the current practice of landscape validation by qualitative analysis requires highly specialized skills and domain knowledge. Furthermore, the computational efficiency of TMELand should be improved in the future. While Python is sometimes less efficient than other languages like Java or C++, we can parallelize the program (e.g. by using GPU) to speed up TMELand.

## Acknowledgements

C.L. is supported by the National Key R&D Program of China (2019YFA0709502) and the National Natural Science Foundation of China (12171102). We would like to thank Mr. Pei Lin for early participation in implementing the TME algorithm in Python.

---

**Lin Zhu** is currently a Master’s student in the School of Information Science and Technology at the ShanghaiTech University, Shanghai, China. She received her B.S. from Chongqing University. Her research direction is machine learning in bioinformatics and systems biology, with a focus on methods for dynamical and datadriven modeling of gene regulatory networkbased Waddington’s epigenetic landscape, as well as the development of related software.

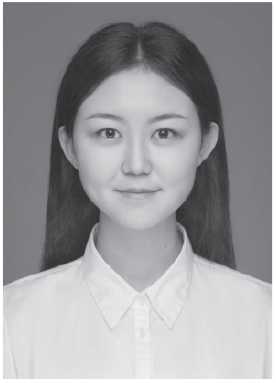

**Xin Kang** received the Ph.D. degree in Applied mathematics from Fudan University, Shanghai, China, in 2021. She received her B.S. from Jilin University. Her research is centered on the stochastic dynamics of gene regulatory networks.

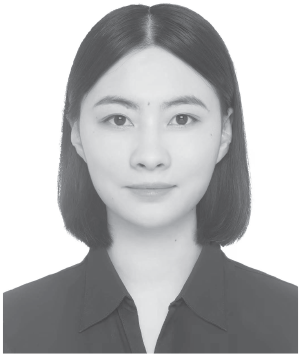

**Chunhe Li** is an Associate Professor in Shanghai Center for Mathematical Sciences at Fudan University. He is also a faculty affiliated with the School of Mathematical Sciences and Institute of Science and Technology for Brain-Inspired Intelligence at Fudan University. Dr. Li received his B.S. from Tianjin University, and Ph.D. in Analytical chemistry from Chinese Academy of Sciences in China. Prior to joining Fudan University in 2016, he worked as a postdoctoral fellow at Stony Brook University and University of California at Irvine in USA. Dr. Li’s research focuses on computational biophysics, systems biology, stochastic dynamics of gene regulatory networks, and data mining. He develops novel approaches for quantifying stochastic dynamics of gene networks to understand the underlying mechanisms of cell fate decisions in specific biological systems, such as cell cycle, development, and cancer.

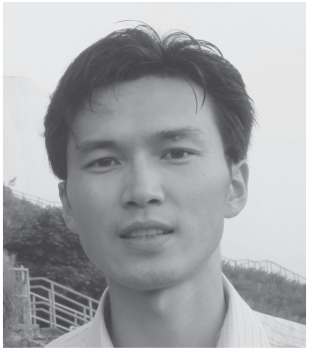

**Jie Zheng** is an Associate Professor (tenured) at the School of Information Science and Technology, ShanghaiTech University, Shanghai, China. He received his B. Eng (honors) from Zhejiang University in China, and his Ph.D. from the University of California, Riverside in USA, both in Computer Science. He worked as a Postdoctoral Visiting Fellow and Research Associate at the National Center for Biotechnology Information (NCBI), National Library of Medicine (NLM), National Institutes of Health (NIH), USA, and as an Assistant Professor at the School of Computer Science and Engineering, Nanyang Technological University (NTU), Singapore. Dr. Zheng’s current research interests are bioinformatics, biomedical data science, AI for drug discovery and precision medicine. He develops novel algorithmic and AI methods (e.g. machine learning and data science techniques, data-driven and knowledge-driven models) to help answer biomedical questions.

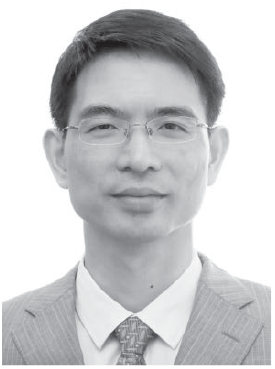

